# Significant non-existence of sequences in genomes and proteomes

**DOI:** 10.1101/2020.06.25.170431

**Authors:** Grigorios Koulouras, Martin C. Frith

## Abstract

Nullomers are minimal-length oligomers absent from a genome or proteome. Although research has shown that artificially synthesized nullomers have deleterious effects, there is still a lack of a strategy for the prioritisation and classification of non-occurring sequences as potentially malicious or benign. In this work, by using Markovian models with multiple-testing correction, we reveal significant absent oligomers which are statistically expected to exist. This strongly suggests that their absence is due to negative selection. We survey genomes and proteomes covering the diversity of life, and find thousands of significant absent sequences. Common significant nullomers are often mono- or dinucleotide tracts, or palindromic. Significant viral nullomers are often restriction sites, and may indicate unknown restriction motifs. Surprisingly, significant mammal genome nullomers are often present, but rare, in other mammals, suggesting that they are suppressed but not completely forbidden. Significant human nullomers are rarely present in human viruses, indicating viral mimicry of the host. More than 1/4 of human proteins are one substitution away from containing a significant nullomer. We provide a web-based, interactive database of significant nullomers across genomes and proteomes.

## Introduction

The terms *minimal absent words* (MAWs) and *nullomers* both describe the shortest sequences that do not occur in the entire genome or proteome of an organism. Although many biotechnological applications have been envisioned, from potential selective drugs (1–2) to forensic practice (3), the actual role of nullomers has intensely been debated (4) and still remains enigmatic. Lately, fast tools and efficient algorithms have been introduced making the discovery of globally missing sequences practical (5–8, 42, 56–57). In 2012, Alileche and colleagues demonstrated that two absent *5*-amino-acid peptides cause fatal damage to cancer cells (2), while *5* years later Alileche and Hampikian showed that the same MAWs have a broad lethal effect on cancer cell lines derived from nine organs represented in the NCI-60 panel (1). Additionally, Silva and colleagues have reported that three minimal *12*-nucleotide fragments entirely absent from the human genome, appear consistently at the same location in two protein-coding genes of Ebola virus genomes (9). Although there exists an unambiguous sequence conservation in their findings, it is not clear whether the absence of these oligomers is statistically expected. In the same study, the term minimal *relative absent words* (RAWs) is introduced, describing sequences that are present in a pathogenic organism but absent from its host. On one hand, all the above findings ideally support the conjecture that MAWs may have gone extinct due to evolutionary pressure or putative deleterious effects. Perhaps the unfavourable properties of nullomers are linked with forbidden spatial conformations followed by functional consequences incompatible with life. Hence, the structural arrangements of globally absent motifs and the putative perturbation of molecules upon their appearance (i.e. emergence of a nullomer upon a mutation) form an interesting area for future research. Conversely, a finite set of sequences, for example the entire genome or proteome of a species, does not include all the different combinations of elements from the alphabet it is composed of, due to the fact that the combination of residues in a sequence increases exponentially with its length. Therefore, it still remains a riddle whether the absence of a nullomer is, in fact, an evolutionary consequence linked with adverse effects or a product of randomness.

In this study, we introduce a robust probabilistic method named *Nullomers Assessor* (https://github.com/gkoulouras/nullomers-assessor) for the evaluation of globally missing sets of oligomers in any species, considering the fact that biological sequences of living organisms are driven by mutational biases and natural selection, and consequently are not entirely random (10). Naturally occurring sequences present patterns and combinatoric properties which can be signatures for the identification of functional elements as such promoters, tandem repeat expansions, introns, exons, and regulatory elements (18). In addition, evolutionarily well-separated species are known to possess distinct statistical characteristics in their DNA or peptide sequence chains (19). All these distinctive properties of biological sequences have frequently been studied using probabilistic models. Markov chain models have been widely and successfully employed in various biological problems including sequence analysis in the past (26–29). Taking advantage of these particular properties of biological sequences, we developed a method which approximates the likelihood of an absent sequence to occur exactly zero times, in order to address the following three questions. First, are there statistically significant minimal absent sequences in biological species; in simple words, what is the expected probability for an actual nullomer to be indeed absent, based on the compositional pattern in the full genome or proteome of a species? Second, are there significant MAWs in common across evolutionarily diverse living organisms? And finally, does the creation of a previously absent sequence perturb a molecule; more precisely, are there mutations with functional or stability impact which at the same time ‘generate’ a nullomer?

Furthermore, given the possible relevance of nullomers to diverse research areas, as for example the under-studied ‘dark’ majority of the genome (11, 12), patterns and evolutionary features between viruses and host species (31–34, 76), rare variants (13–15) as well as cancer driver mutations in non-coding regions (25), we provide the community with the results of the present study in the form of a publicly-available, downloadable and web-accessible repository of significant absent motifs, named *Nullomers Database* (https://www.nullomers.org/). Modern web-technologies and visualization features have been harmonically combined resulting in a dynamic and user-friendly environment. To the best of our knowledge, this is the first attempt for a centralised, open-access and searchable resource of non-occurring genomic and peptide sequences. *Nullomers Database* is intent on being a periodically updated and continuously enriched repository of significant absent sequences from various organisms, as they have been assessed by *Nullomers Assessor*. We are hopeful that the intuitive and interactional graphical user interface of *Nullomers Database*, in conjunction with the integrated annotation and the powerful searching features that it employs, will facilitate exploration and shed new light on the puzzling and, up to the present time, little-known world of nullomers.

## Materials and Methods

### Identification of Minimal Absent Words

The identification of nullomers was achieved using the MAW console application (5), an open source 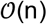-time and 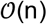-space algorithm for finding minimal absent words based on suffix arrays. It is fast and straightforward for genomes and proteomes of moderate length, because all calculations are performed in-memory. On the other hand, when applied to long sequences of size *n* the algorithm requires more than 20 ∗ *n* bytes of available RAM, which causes a bottleneck on large datasets such as the human genome. For the detection of nullomers on sizable datasets we used the em-MAW software tool (6), a marginally slower alternative which utilises external memory. Both MAW and em-MAW require an input fasta file which contains the whole genome or proteome of an organism, as well as two numerical arguments that indicate the shortest and longest nullomers to search for. Throughout the study, we searched for sequences of length between 4 and 14 nucleotides which are globally absent from both forward and reverse-complement strands. For peptide sequences we set the identification range between minimum 4 and maximum 6 amino acids in order to keep complexity at a reasonable level. The output of both MAW and em-MAW is a list of missing sequences of a given dataset.

In our analysis, we downloaded full proteomes of two main organisms (i.e. *Homo sapiens* and *Mus musculus*) from UniProt (30; https://www.uniprot.org/proteomes/), while a range of more than 1.500 genomes from archaea, bacteria, protozoa, fungi, invertebrates and vertebrates were retrieved via NCBI Genome (35; ftp://ftp.ncbi.nlm.nih.gov/genomes/). We developed custom Python scripts to discard headers and concatenate sequences from multiple fasta files, to produce files with one header and one-line sequence for each organism. The above step was applied to protein sequences as well, because MAW and em-MAW are developed in a way to calculate nullomers of each individual record in a fasta file. For the identification of peptide nullomers, we combined information both from the Swiss-Prot and TrEMBL sections of UniProtKB including protein isoforms. The redundant but slightly different sequences (isoforms) that were included in this step can contribute to the elimination of incorrectly identified MAWs. Moreover, the incorporation of predicted with manually reviewed records produces a list of more confident non-occurring sequences. The final pre-processing step included the removal of any ambiguous residues (character N in genomes or B, J, X, Z in proteins). Eventually, we generated lists of global nullomers from various species which were used as an input for downstream analysis in *Nullomers Assessor*.

### Applying Markov models to genomic and protein sequences

Statistical models represent the observed variability in data by probability distributions. A simple model of a sequence *X* = (*X*1, *X*2, *X*3, …) is a first-order Markov chain, where each position is dependent only on its immediate precursor. For example, the probability of observing a ‘G’ at one position depends (only) on whether there is a ‘C’ in the previous position. This can be expressed as:

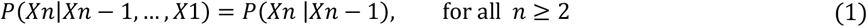

In our method, we consider genomes and proteomes (hereinafter background sequence) as Markov chains. A first order Markov chain is a model where each position is contingent merely on its previous position. Likewise, in an *n*-th order Markov chain, each position depends on the *n* previous positions. In order to decide whether a nullomer is statistically expected to exist, we estimate Markov probabilities. First, the frequencies of elements (nucleotides or amino acids) of the background sequence are calculated. Then, three Markov probability matrices are generated, one for each of the first three Markov model orders. In general, a substitution matrix for *m* distinct letters (e.g. *m* = 4 for DNA) and order *n*, is a grid of *m* ∗ *m*^*n*^ probabilities. As an example, a stochastic matrix of third-order for all the naturally occurring amino acid residues in a bacterium (20 distinct amino acids assuming that neither selenocysteine nor pyrrolysine are present) requires a matrix of 20 ∗ 20^3^ = 160.000 cells (namely 20 rows and 20^3^ columns, or vice versa). Each of these 160.000 probabilities indicate the likelihood for a specific amino acid to occur given the three preceding amino acids.

In non-mathematical terms, each residue of a biological sequence is dependent on the *n* previous elements, where *n* defines the order of a Markovian process. Implicitly, a stochastic process in a biological sequence can reveal the sequential preferences among neighboring residues as well as reflect avoided motifs which may introduce an unfavorable structural folding. We utilised the above-described fundamental mathematical notion and developed a custom Python script (*Nullomers Assessor*) which approximates the likelihood for a nullomer to occur zero times based on the first four orders of Markovian chains (including zeroth order). More precisely, four distinct *p-*values are assigned to each nullomer of the list. To elucidate and put this probabilistic property into a more biological context, we provide the following example. Assuming the peptide ‘PTILA’ is an absent minimal 5-mer, then the probability of being entirely absent based on a second-order Markov chain can be calculated as illustrated below:

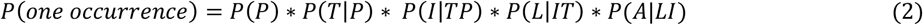

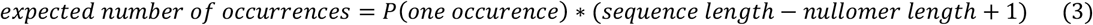

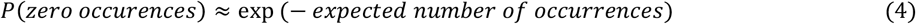

Initially, the probability of ‘PTILA’ to occur is estimated (Equation 2). In this formula, P(P) denotes the observed frequency of proline in the entire background sequence. Subsequently, P(T|P) signifies the probability of threonine to arise after a proline. Similarly, P(I|TP) indicates the probability of isoleucine to occur given the previous 2 adjacent amino acids are a proline followed by a threonine, and so forth. Next, the expected number of occurrences for the specific actual nullomer is estimated (Equation 3), a value that is used for the final estimation of the second order zero-occurrence probability (Equation 4). In a similar manner, the probabilities of first and third orders are computed, while the zeroth order simply mirrors the frequency of residues in the background sequence. In simple words, the calculated *p*-values represent the probability of a minimal absent word to be indeed absent based on the rate at which the residues occur as well as three additional transition probabilities which reflect the frequency of 2-, 3-, and 4-mers of the examined genome or proteome.

Although 4 different probabilities are calculated, the maximum p-value is kept and assigned to each examined nullomer. Since a *p*-value denotes the chance of a nullomer to occur exactly zero times (namely to be absent), then the lower the value, the more expected the nullomer is to exist (or equivalently the less expected it is to be, indeed, absent). By keeping the highest probability amongst the 4 calculated p-values, we expect to end up with fewer but more confidently true-positive results.

### Multiple hypothesis testing and statistical correction

There is a large number of short sequences that each have a chance of being absent. Therefore, the emergence of false positive results must be controlled (46). More specifically, all the 4 calculated *p*-values of each nullomer are corrected for Type I errors and readjusted based on one of three statistical correction methods which are provided built-in with the current version of the tool. In exact terms, users can choose between Bonferroni (47), Benjamini-Hochberg (48; widely known as False Discovery Rate or simply FDR), or Tarone method (49) which is a modified Bonferroni procedure. The Bonferroni correction is particularly conservative, especially when applied to proteome datasets, due to the large alphabet size and high number of tests. More specifically, each of the 4 individual *p*-values is multiplied by the count of all possible different *k*-mers of length *k*. For example, the corrected *p*-value of a genomic *k*-mer is the product of the actual *p*-value multiplied by 4^*k*^, where *k* denotes the length of the examined absent motif. For peptide nullomers the multiplier changes to 20^*k*^. Generally, the following formula illustrates the Bonferroni correction step:

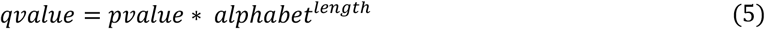

Alternatively, in the Benjamini-Hochberg procedure, the probabilities are sorted in descending order and sequentially rejected if the product of a *p*-value and the number of remaining tests is greater than a cut-off limit. The FDR method though, which constitutes a milder alternative, performs markedly loosely when applied to large eukaryotic genomes resulting in thousands of significant nullomers. This motivated us to incorporate a third correction option, the Tarone methodology. This is a special case of the Bonferroni method, where we make it less conservative by doing fewer tests. For a given word-length *k*, we calculate the zero-occurrence probability (the maximum of the four Markov probabilities) for each of the 4^*k*^ *k*-mers. Then, these 4^*k*^ *k*-mers are ordered in descending order of zero-occurrence probability. Next, we exclude from testing any *k*-mer whose absence would not be significant because the Bonferroni-adjusted p-value is above a cut-off threshold:

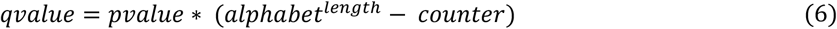

Equation (6) represents the mathematical notation of the Tarone method, where *counter* denotes a number which progressively increases by 1 when a *k*-mer is excluded from testing. Finally, the “testable” *k*-mers that remain are compared to the actual nullomers in the list, and those in the intersection are output. In this way, the stringent nature of Bonferroni remains, whilst a milder adjustment is performed every time a test is excluded due to impossibility of significance.

Next, we tested the accuracy and rigour of the 3 different correction methods using a common dataset and a constant threshold of false discovery control. The results show that Bonferroni correction performs the strictest cleansing of false positive nullomers (very likely at the cost of increased false negatives), followed by the Tarone method, whereas the FDR approach constitutes the least stringent alternative. More precisely, any result-set derived by Bonferroni is a subset of the corresponding Tarone set, while the results of the FDR method almost always include all the above outcomes. Throughout our analysis, a fixed false discovery threshold of 1% has been applied, both when searching for genomic or peptide nullomers. In order to eliminate the emergence of Type I errors to the utmost degree, we report a nullomer as significant only when all four corrected probabilities are lower than the user specified cut-off. However, we advise users to apply Bonferroni correction to relatively large genomes, in order to control as much as possible the emergence of false-positive results, whilst starting with an alternative method when dealing with either small genomes (i.e. viruses or microbe datasets) or proteomes in general.

### Testing precision by shuffling input sequences

Random sequence shuffling is a widely used approach to evaluate stochasticity as well as statistical significance of results. In order to evaluate the rigour of our method, especially because Equation (4) is only approximately true, we performed permutation tests by randomly shuffling the human proteome. Since *Nullomers Assessor* calculates up to third-order probabilities, we sought to retain unaltered not only the counts of distinct amino acids, but also higher-order statistics, such as the frequency of adjacent letters (doublets and triplets) of the entire proteome. For this purpose, we used a sophisticated shuffling algorithm, uShuffle (21), which performs random shuffling of sequences while preserving *k*-let counts. The C# software package of the uShuffle method was used to shuffle the human proteome 10 times while preserving the frequency of amino acids, doublet occurrences and tripeptides. Next, for each of the 10 shuffled proteomes we re-generated lists of nullomers in order to examine whether any of the new random absent sequences would come into view as significant. By keeping the singlet, doublet, and triplet amino acid frequencies unchanged but not their order, we expected to end up with utterly different lists of nullomers. We used the original human proteome as a background sequence and re-ran our method 10 times (once per each new list of nullomers) in order to assess the newly created lists of ‘counterfeit’ absent words. Even though the script was executed using identical parameters (background proteome, correction method, threshold of statistical significance), no significant results emerged in any of the 10 attempts, demonstrating the stringency of our methodology. Thus, we assessed 10 sets of random missing sequences using the real transition probabilities of the reference human proteome given the fact our method scans any background sequence by considering frequencies up to 4-mers (precisely up to third-order Markov chains). The outcome of this step strongly suggests that *Nullomers Assessor* is able to disclose truly significant nullomers.

## Results and Discussion

In the present study, we rigorously assess absent sequences for their statistical significance. We examine genomes and proteomes from hundreds of organisms (Table 1) and show lists of nullomers which are unexpected to be absent, in contrast with other missing sequences. Our findings demonstrate that several thousands of absent sequences are statistically expected to occur in various genomes. The longest significant human genomic MAW is composed of 13 nucleotides, whilst the lengthiest significant peptide nullomer from the same organism is 5 residues long. After applying our method to the entire human genome, 13 significant genomic nullomers stood out (Bonferroni correction at 1% cut-off) from a set of more than 27 million non-occurring oligomers. In essence, the specific 13 missing words are highly statistically foreseen to occur somewhere in the human genome but, in reality, they are totally absent. In a similar manner, we analysed peptide nullomers from the human and mouse proteomes. Thirteen absent peptides from the human proteome were classified as significant when the Tarone correction method was used, while eight peptides emerged when we applied identical parameters to the mouse proteome (Table 3). Moving the hypothesis of harmfulness a step forward, we systematically explored nullomer-making mutations which are one residue away from the reference sequence. More specifically, we calculated all the possible single amino acid substitutions in all protein records of each proteome that can give rise to any of the total 21 significant absent words. This might offer useful insights for unravelling plausible mechanisms of evolvability that underlie peptide nullomers. Prior research suggests that different residues differ in respect of their mutational preference (41) and reports implications in phosphorylation sites (37). Therefore, the mutational landscape of nullomers presented in this work, may provide a useful resource for future sequencing studies, especially in the field of proteomics (40). To this end, we highlight and share more than 30.000 candidate nullomer-making alterations in the form of interactive visual components via *Nullomers Database* web-portal. In addition, we make available a complimentary catalogue of 176 phosphorylation sites (Supplementary Table 1) which can lose their phosphorylation ability upon a mutation and, simultaneously, generate a MAW. Next, we show that the most frequent significant absent words in viral sequences are restriction recognition sites indicating that viruses have probably got rid of these motifs to facilitate invasion of bacterial hosts. It is worth noting that not every species or virus has significant MAWs, thus the provided result-set can be used to reduce the vast nullomers’ space and prioritise non-occurring motifs for future research questions. Finally, we share lists of human nullomers which are seldomly present in viruses suggesting molecular mimicry between virus and host (Supplementary Table 2).

**Table 1.** Summary table outlines the number of analysed genomes and the count of identified significant nullomers per division

| Division | Number of species | Number of total significant nullomers | Number of unique nullomers |  |
| --- | --- | --- | --- | --- |
| archaea | 144 | 2,419 | 1,074 |  |
| bacteria | 559 | 20,082 | 10,547 |  |
| fungi | 14 | 160 | 159 |  |
| invertebrate | 19 | 857 | 711 |  |
| plant | 52 | 4,047 | 3,980 |  |
| protozoa | 13 | 104 | 102 |  |
| vertebrate mammalian | 43 | 471 | 451 | 2,842 |
| vertebrate other | 56 | 3,032 | 2,416 |  |
|  | 900 | 31,172 | 19,440 | 19,415 |

### Genomic nullomers across evolution

To assess whether nullomers have an ancestral origin, we examined a plethora of diverse organisms ranging from bacteria to human. To this end, we hypothesized that distantly-related genomes would share fewer similar sequence features and therefore one would not be surprised to find fewer or no nullomers in common. In contrast, despite the stringent filtering criteria of our methodology, we would expect to end up with identical significant nullomers from closely-related organisms due to the existing high similarity both in genomic and protein sequences. Surprisingly, while some significant nullomers are sporadically shared by some mammals, most are not shared by closely-related mammals (Figure 1). Although none of the 13 genomic nullomers in *Homo sapiens* have emerged significant in *Pan troglodytes*, despite their genomes being ~98% identical, the latter shares 3 absent motifs with the closely related species *Gorilla gorilla* and *Pan paniscus*. Furthermore, *Pan paniscus* shares 2 nullomers with *Saimiri boliviensis* forming a cluster of related organisms with significant results in common. This led us to investigate whether significant absent sequences in human are present in chimp, and vice versa. Since the two genomes are very similar, it might be considered not surprising that absent words in human are rare in chimp. For this reason, we also investigated two more distant species, *Mus musculus* and *Canis lupus familiaris*. Then, for each significant nullomer of a species, we computed the observed as well as the expected number of occurrences in the other 3 organisms by exploiting again equations (2) and (3) in the *Materials and Methods* section. We found that the majority of nullomers in *Homo sapiens* is present in *Pan troglodytes* (and vice versa) with a median frequency of 1 occurrence, while the median number of expected occurrences of the significant human MAWs in chimp is 47 (Figure 2A). In a like manner, the estimated median frequency of chimp-derived nullomers in the human genome is 53 (Figure 2B) while the detailed dataset is provided in Supplementary Table 3. Figure 2C and 2D outline a similar trend in nullomers of *Mus musculus* and *Canis lupus familiaris*, respectively, where the expected number of occurrences is again more than the actual observations. This finding suggests an alternative hypothesis in which significant nullomers are not completely forbidden, but they are strongly suppressed. Strongly suppressed sequences are expected to occur just a few times, so by chance fluctuations some of them could appear zero times.

**Figure 1.**
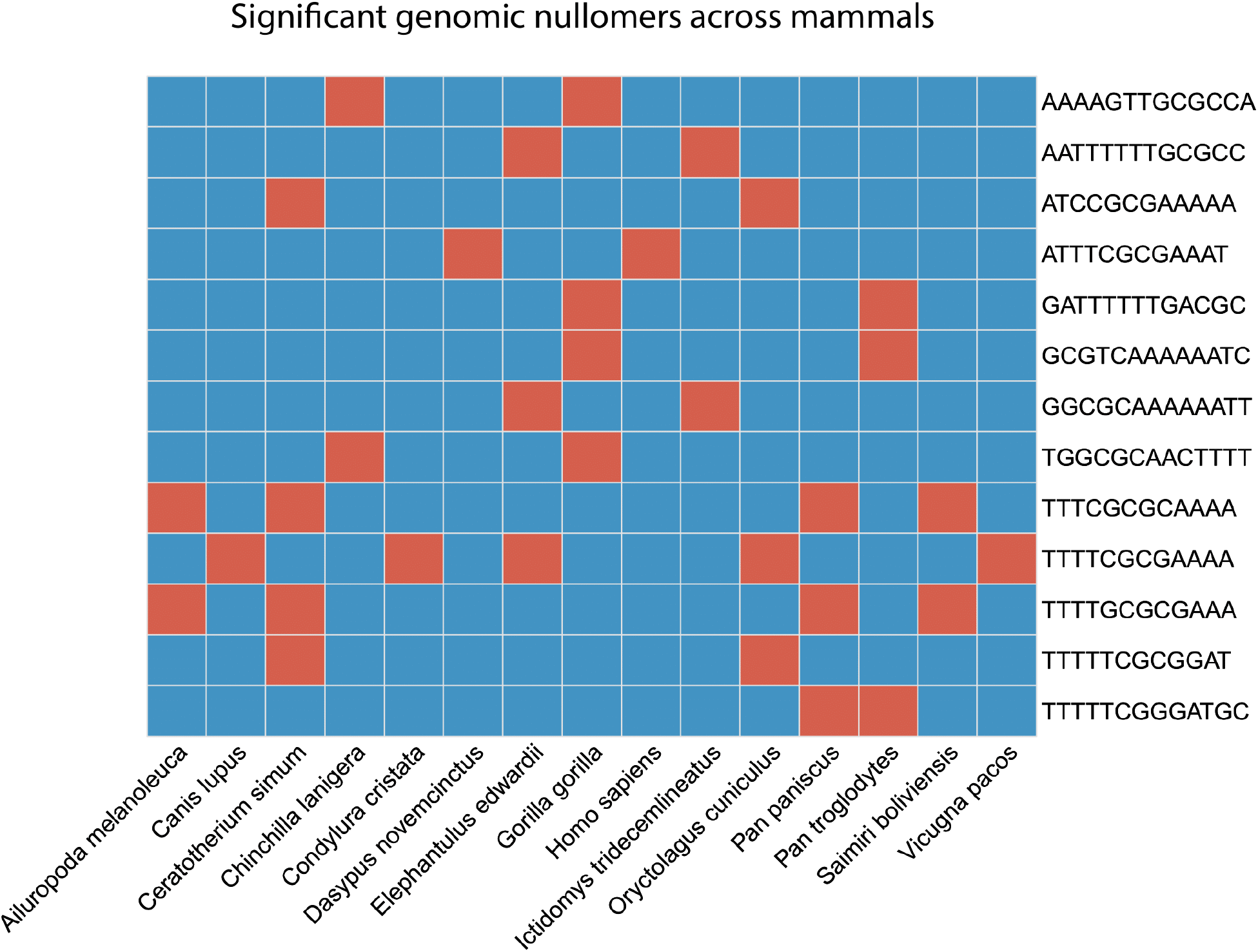
Comparison of significant genomic absent sequences across mammals. Only nullomers that are shared in at least two species are shown. A red grid-cell indicates a significant nullomer (evaluated by *Nullomers Assessor)* while blue colour denotes a non-significant or present motif.

**Figure 2.**
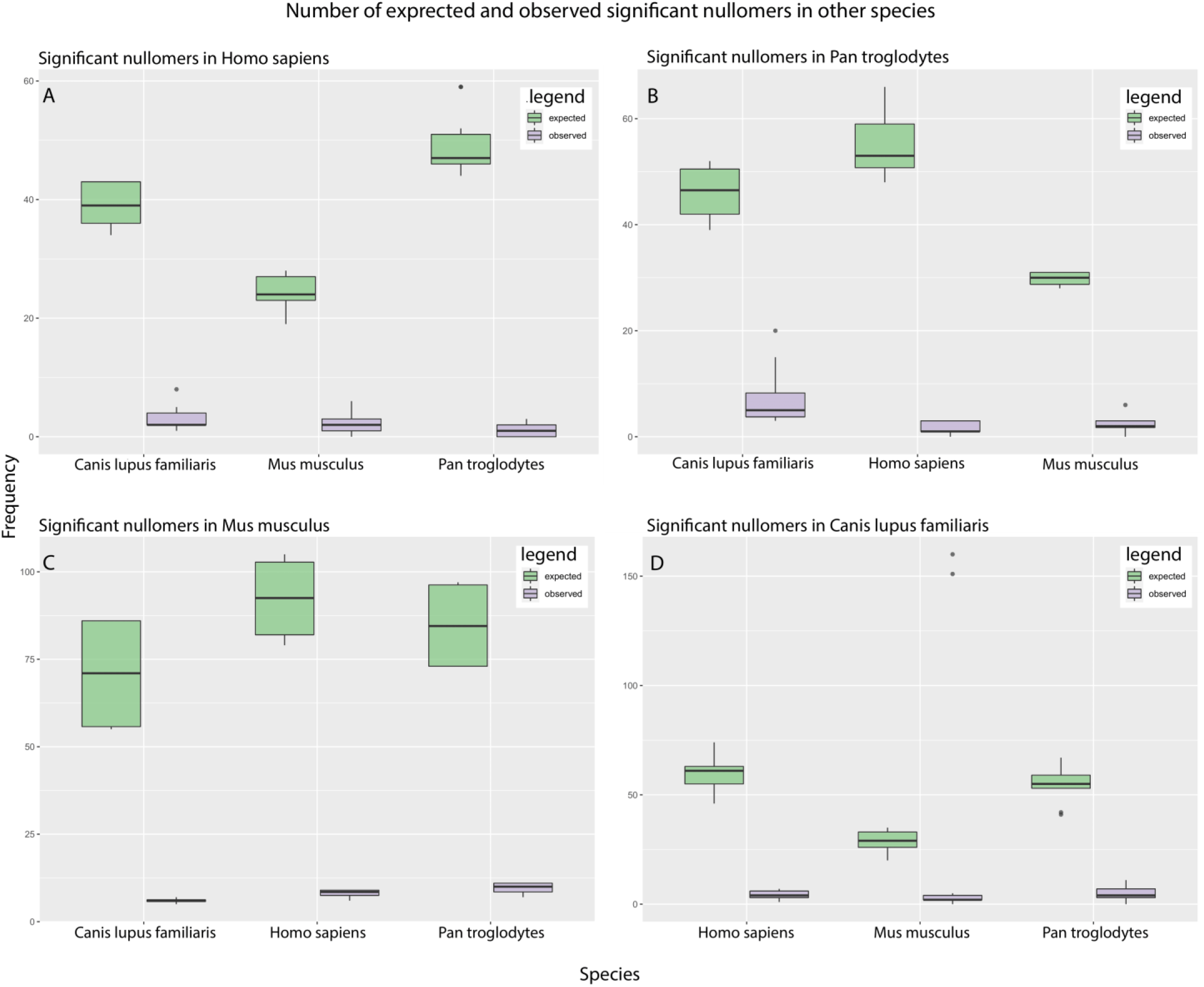
(A) Comparative quantitative analysis of human-derived genomic nullomers that exist in Pan troglodytes, Mus musculus and Canis lupus familiaris. The green boxplots show the expected number of the total nullomers in Homo sapiens (13 in total) while the purple boxes correspond to the observed frequencies. Similarly in (B) the nullomers of chimpanzee have been search against the genomes of Homo sapiens, Mus musculus and Canis lupus familiaris. In (C) and (D) the nullomers of Mus musculus and Canis lupus familiaris, respectively, have been searched against the other 3 species.

In contrast, none of the roughly 2.850 unique significant nullomers in vertebrates are shared with any of the other species which belong to archaea, bacteria, protozoa, or fungi (Figure 3A). To some extent, this may be due to the heterogenous complexity among species of different kingdoms because significant nullomers of vertebrates are usually longer (Figure 3B). Overall, these observations are in accordance with findings reported by Acquisti et al. (4). The authors of that study have demonstrated that species with a more-recent common ancestor share more MAWs in common in comparison with more distantly-related organisms. Here, we provide evidence that more-closely related organisms share not only common deficits in general, but also some identical significant missing patterns, supporting the hypothesis of evolutionarily-conserved aversion to these sequences.

**Figure 3.**
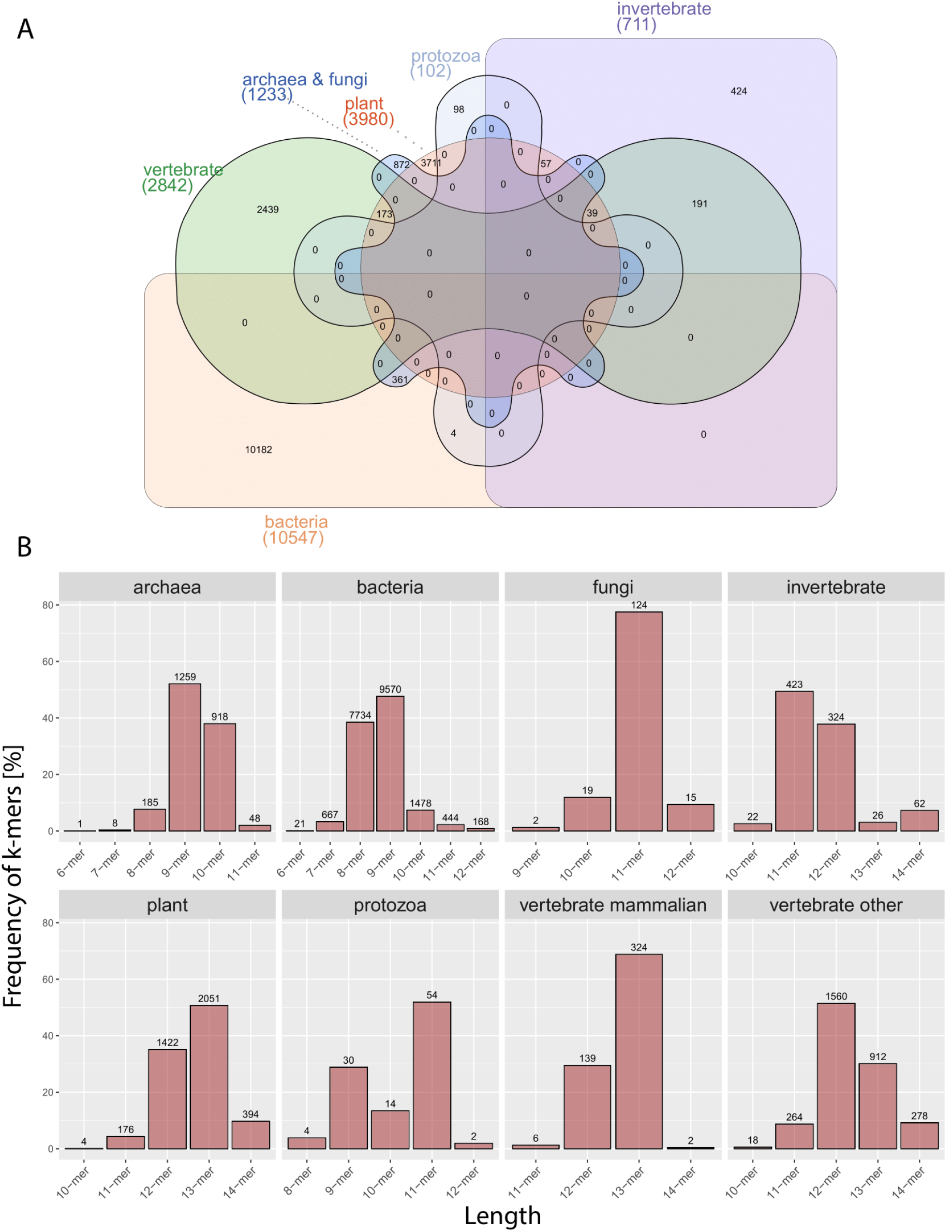
(A) Venn diagram showing the number of shared nullomers derived from 900 genomes grouped by division. Created using InteractiVenn (74; http://www.interactivenn.net/index2.html) (B) Distribution of k-mers length per division. Each bar represents the count of nullomers for a specific motif length.

Furthermore, we observed multiple genomic MAWs in common across species ranging from 6 to 14 residues. Table 2 presents the most frequent significant nucleotide sequences whose absence is shared between at least 18 species, while a complete list is in Supplementary Table 4. Thus, identical significant nullomers which are shared across species might be usable for biotechnological applications or further research. What is noticeable in this dataset is the high rate of mononucleotide tracts. It has previously been shown that enrichment of poly(A) tracts is linked with important functional roles, including DNA methylation (62), ribosome stalling (63), and translational efficiency (64), to name a few. The longest poly(A) sequence in our dataset is a 11-nucleotide poly(A) motif absent in 18 bacterial and archaeal species, while a 12-mer poly(T) tract has been marked significant in *Clostridium sartagoforme*, implying the high content of repeated adenine and thymine in their genomes, respectively. In *Homo sapiens*, the ‘ATTTCGCGAAAT’ nullomer constitutes one of the nearly 250 palindrome nullomers of the result set and it is shared with *Dasypus novemcinctus* and *Gekko japonicus*. Table 4 details the frequency of mononucleotide tracts per length, while all the palindromic nullomers can be effortlessly accessed and downloaded via *Nullomers Database*.

**Table 2.**
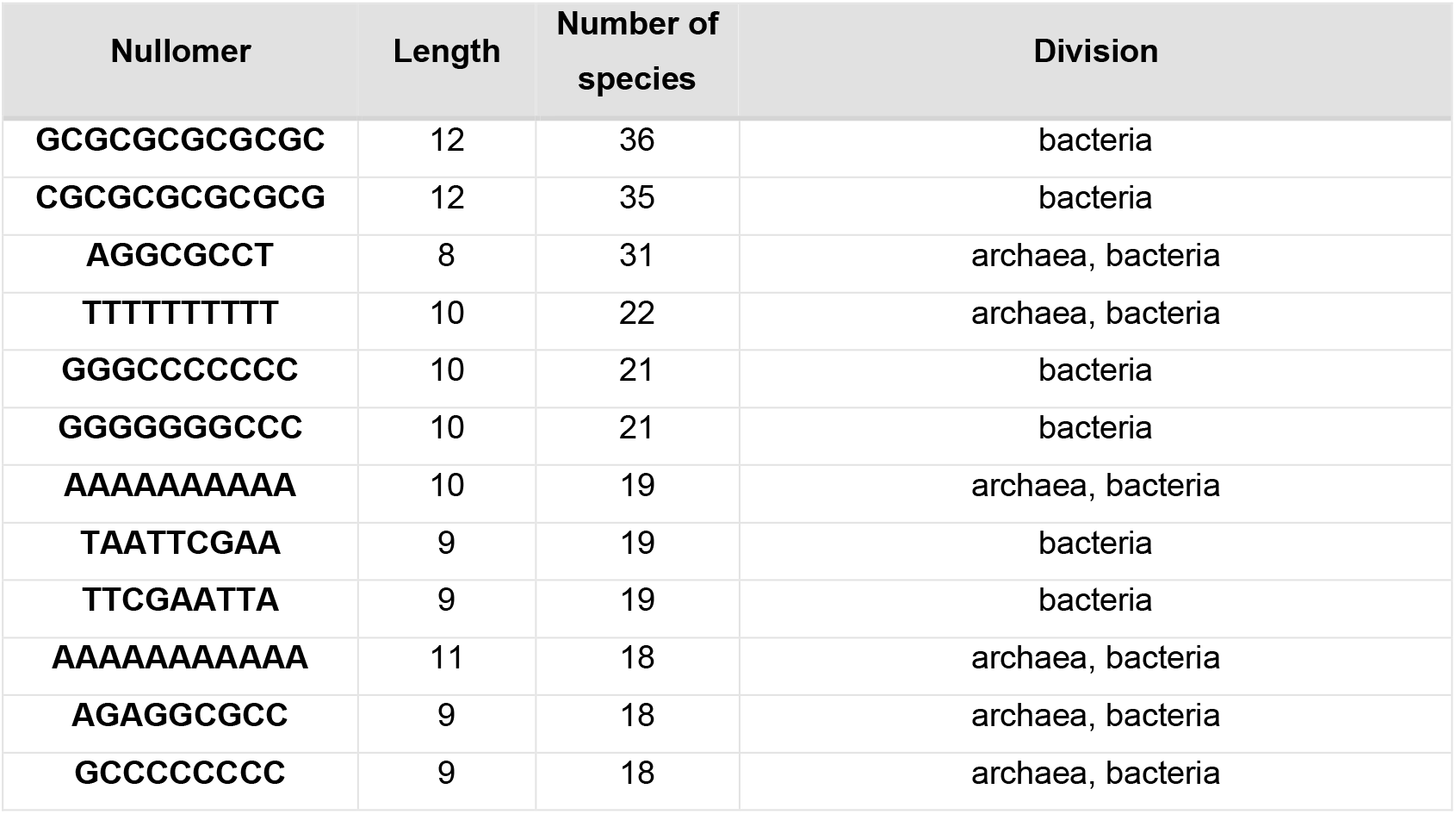
List of the most frequent significant nullomers across species.

### Peptide nullomers and nullomer-making mutations

According to the definition, a nullomer is a minimal-length sequence that does not occur within a longer sequence. Consequently, if we think of a *k* − 1 peptide derived from a nullomer of length *k* without either the first or the last amino acid, then the remaining part of *k* − 1 residues certainly occurs somewhere in the background sequence. This information is enough to identify regions in proteins that are susceptible to create a significant nullomer upon a mutation. Additionally, single amino acid alterations capable of generating a nullomer can occur in non-edge positions of a k-mer. To illustrate this point further, we provide the following example. BRCA1 is a human gene that produces a tumour suppressor protein of 1863 amino acids (UniProt ID - P38398) involved in various biological mechanisms including DNA damage repair and embryonic development (38,39). Two possible nullomer-making mutations Ser > Pro and Lys > Asn in positions 628 and 1171 can give rise to the ‘LVVPR’ and ‘INESS’ absent words, respectively. Therefore, the actual amino acid chains ‘LVVSR’ and ‘IKESS’ that normally exist in the reference protein sequence could be considered ‘sensitive’ to putative S628P and K1171N mutations in the 4^th^ and 2^nd^ position, respectively. Moving this simple idea forward in conjunction with the scenario of noxiousness behind entirely missing sequences, we hypothesised that nullomer-making mutations would potentially introduce unfavourable effects to these molecules. To this direction, we developed an automated procedure that detects positions in proteins which are susceptible to generate one of the significant identified nullomers upon a single amino acid alteration. We applied this to the entire *Homo sapiens* and *Mus musculus* proteomes ending up with a list of 34.053 positions which are prone to create one of the 21 significant nullomers in both organisms (Table 3). In the human proteome, 21. 668 potential alterations can lead to one of the 13 significant minimal absent peptides in 16.045 UniProt IDs (of which 6.576 belong to unique manually reviewed records). This suggests that more than one fourth of the human proteome is susceptible to introducing an utterly absent sequence, with a single amino acid alteration. With this information in hand, we investigated evolutionary tendencies of mutability (amino acids to be mutated) and targetability (resulting amino acids upon a mutation) in nullomer-making positions (Figure 4, 5). A clear propensity is apparent in targeted valine in the human proteome, while the extremely high number of mutations to isoleucine in both organisms constitutes an intriguing observation considering that Ile is one of the rare amino acids (frequency <5%). At the opposite extreme, 5 amino acids (cysteine, histidine, methionine, tryptophan, tyrosine) are null targets of human nullomer-making mutations, a reasonable fact given that none of the 13 significant nullomers have any of these amino acids in their sequences. Similarly, asparagine, cysteine, histidine, methionine, phenylalanine, tryptophan, and tyrosine are not nullomer-making target amino acids in Mus musculus. We produced matrices of mutational transitions in order to further detect possible tendencies (Figure 5). The resulting plots demonstrate that Leu > Val and Leu > Ile are two prevalent alterations in human, whereas Ala > Ile and Glu > Ile are the most common nullomer-creating substitutions in the mouse proteome. Curiously, Leu, Ile, and Val are precisely the three branched-chain amino acids. Next, we investigated whether the inclusion of non-reviewed proteins (records from TrEMBL) affects the nullomers’ mutational space. What can be seen in Figure 5 is that dominant mutational trends remain unaffected either with or without predicted records.

**Table 3.**
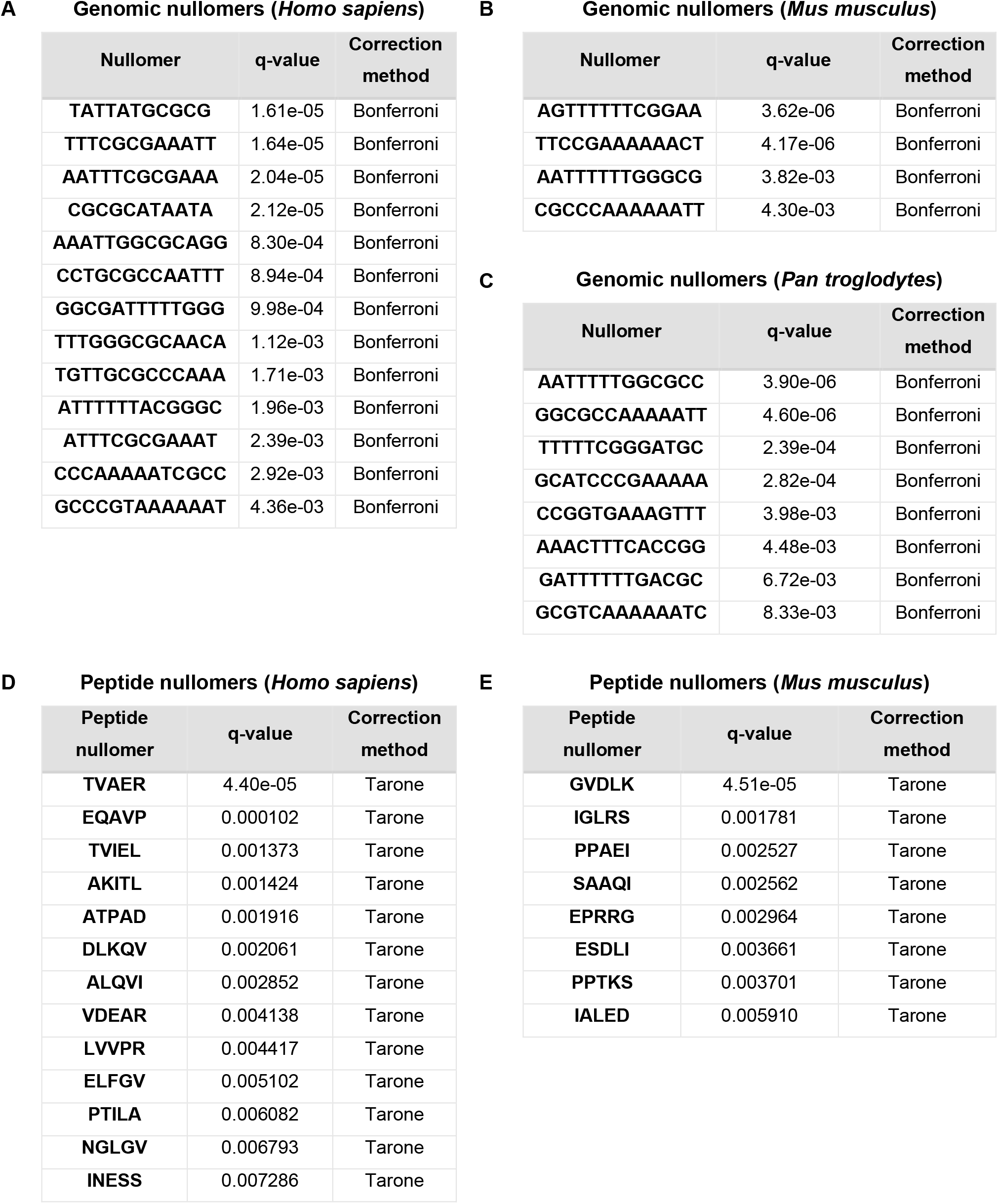
List of significant genomic nullomers in (A) Homo sapiens, (B) Mus musculus and (C) Pan Troglodytes. List of significant peptide nullomers in (D) Homo sapiens and (E) Mus musculus.

**Table 4.** Summary table of poly-mononucleotide tracts per nullomers’ length

| k-mer length | Count of poly-A tracts | Count of poly-C tracts | Count of poly-G tracts | Count of poly-T tracts |
| --- | --- | --- | --- | --- |
| 9-mer | 79 | 10 | 10 | 93 |
| 10-mer | 46 | 4 | 3 | 63 |
| 11-mer | 18 | 1 | 0 | 19 |
| 12-mer | 0 | 0 | 0 | 1 |
| 13-mer | 0 | 0 | 0 | 0 |

**Figure 4.**
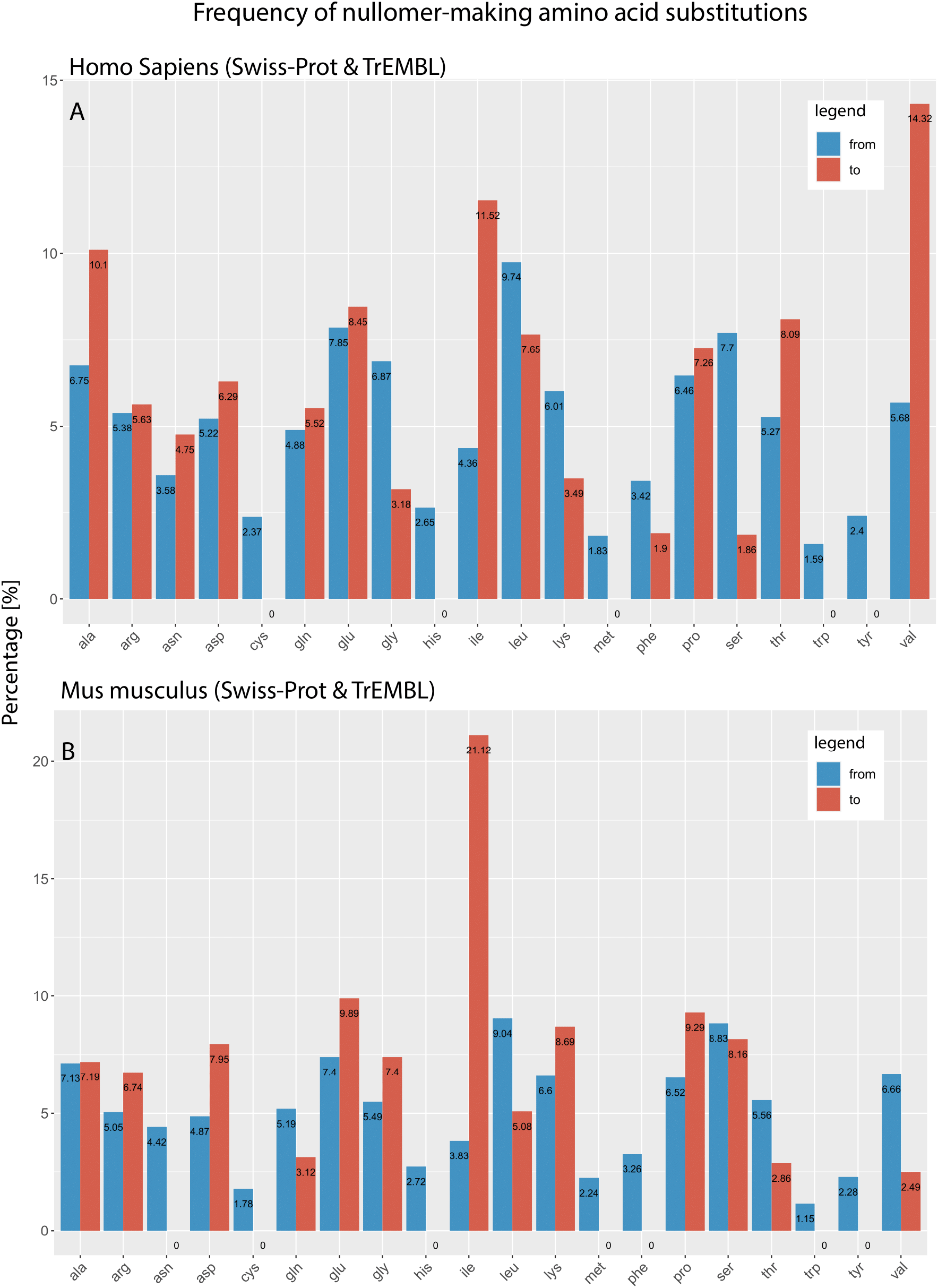
Proteome-wide analysis of all possible nullomer-making mutations in (A) *Homo Sapiens* and (B) *Mus Musculus*. Each bar depicts the frequency of single amino acid alterations which generate a significant nullomer. Blue bars represent the count of mutable amino acids that are prone to alter while red-coloured bars indicate the number of putative nullomer-making substitutions. Proteomes include both canonical, isoform, reviewed and predicted records. Full proteome datasets retrieved from UniProt.

**Figure 5.**
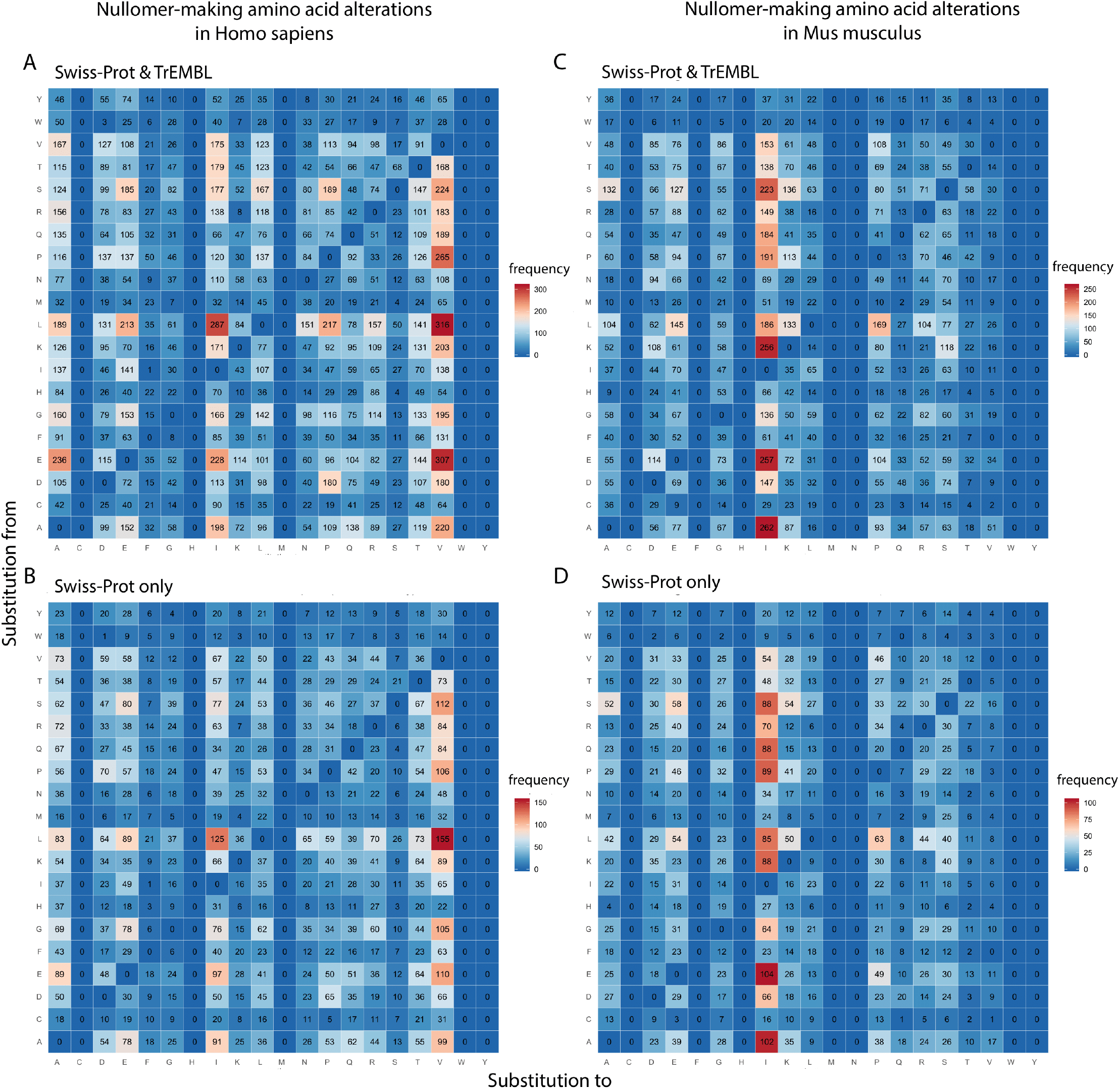
Proteome-wide mutational landscape of nullomer-making alterations. Each grid-cell indicates the frequency of nullomer-making alterations from one amino acid to another. The mutable amino acids are shown vertically, while the target amino acids are depicted in the horizontal axis. Subplot (A) presents nullomer-making substitutions in human protein entries derived from Swiss-Prot and TrEMBL, while only curated records have been considered in (B). Similarly, (C) and (D) depict an identical analysis in Mus Musculus considering reviewed and predicted compared to curated-only records, respectively.

Recently, massive efforts have been put forth to prioritize the functional importance of phosphorylation sites (54,55) as well as decipher correlation between mutation and phosphorylation in cancer (75). For this reason, we compiled a curated list of susceptible nullomer-making phosphosites retrieved from PhosphoSite Plus database (36). Our analysis revealed 176 phosphorylation sites of high confidence which are prone to give rise to a nullomer (Supplementary Table 1), whilst a similar tendency is notable where valine is the most prominent mutational target (Figure 6). Future research should be devoted to characterizing the impact of nullomer-making mutations, especially when a substitution occurs in post translational modification (PTM) sites.

**Figure 6.**
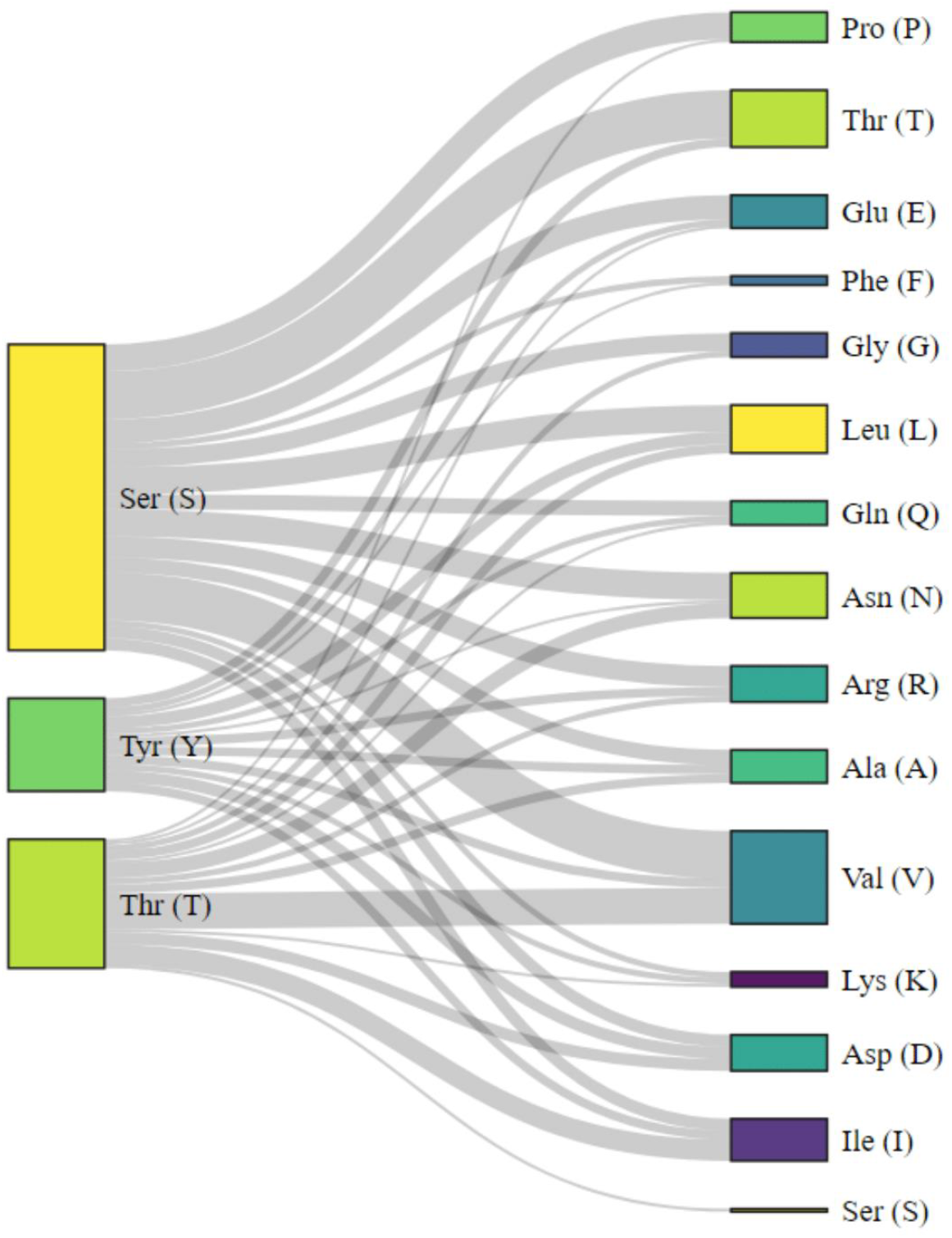
Sankey diagram illustrates nullomer-making mutational trends in human phosphorylation sites. Experimentally verified protein phosphorylation sites have been extracted from PhosphoSitePlus® (v6.5.7) and subsequently matched with nullomer making mutations from Nullomers Database. The left-hand side residues are phosphosites susceptible to alter, while the targeted amino acids are presented in the right side. Box size proportionally indicates the number of putative alterations.

### Relative absent words and nullomers in viruses

Recent studies have shown that viral sequences mimic those of their hosts, to some extent, in part to evade immune responses (32,50–51), and this can be used to predict viral hosts (33). To this end, we set out to survey virus genomes and proteins and investigate whether the scenario of RAWs fits with our previous human-derived findings. Viral sequences and complementary annotation were downloaded from NCBI Virus (43) in March 2020. We filtered out incomplete sequences, whilst we further discarded records that have been isolated from non-human species, resulting to 33.610 genomes and 233.178 proteins. Next, we queried whether significant human nullomers appear in any of the viral sequences. We observed that 5 genomic nullomers (out of the total 13) were present in 39 unique sequences of 3 distinct virus families (Figure 7A). What stands out is the remarkably low number of significant *human*-RAWs which are present in the virus genomes (Supplementary Table 2). In other words, there is a general absence of absent human sequences that are present in viruses.

**Figure 7.**
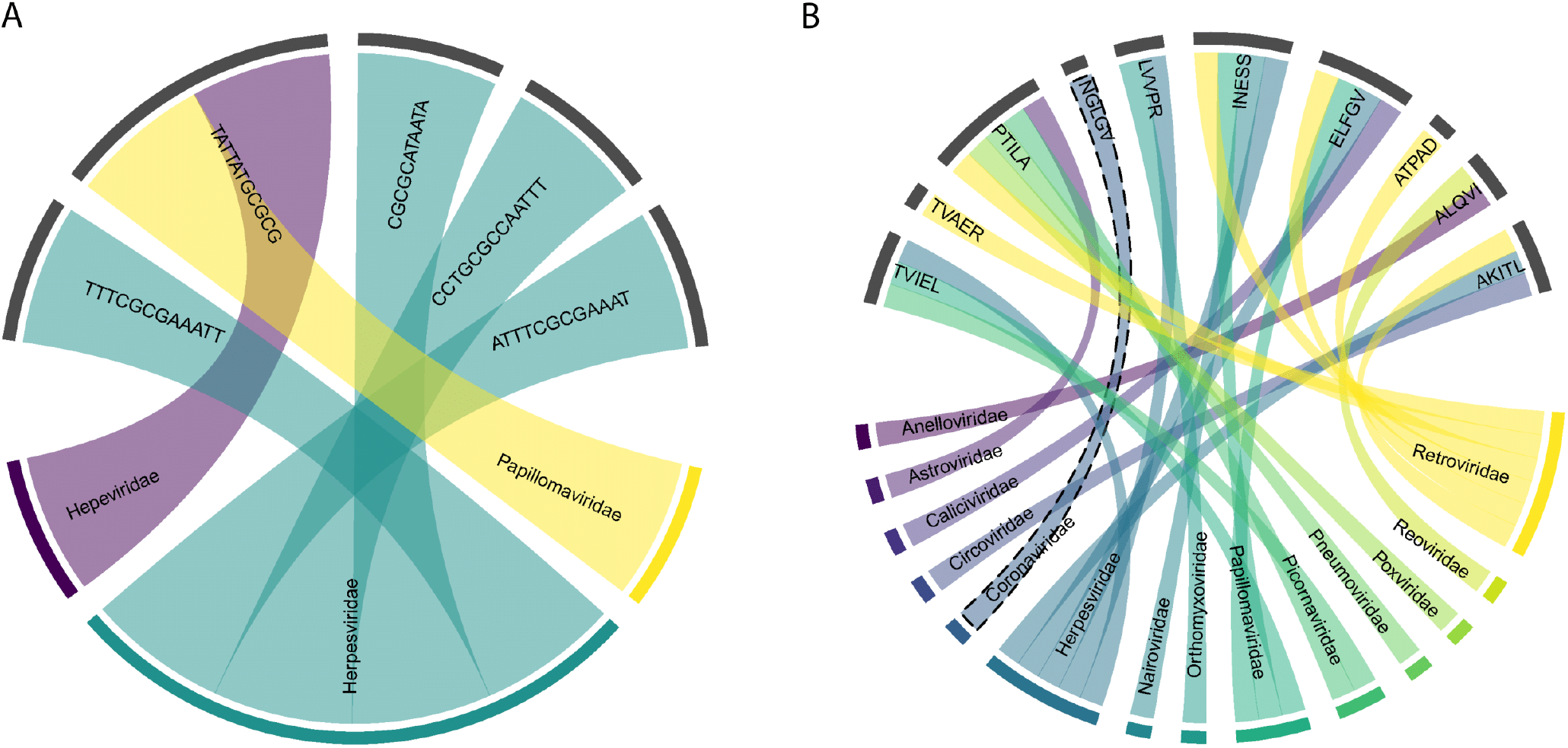
(A) Chord diagram presents 5 significant relative absent words, present in viral genomic sequences (virus families) but absent from the human genome (B) Chord-diagram of correlations between human-derived peptide nullomers and virus families. The highlighted *NGLGV* nullomer has been found conserved in sequences of Coronaviridae family.

Likewise, we investigated whether human-derived peptide nullomers are present in viral protein sequences. We observed that 10 out of the 13 significant human MAWs emerge in various proteins of 14 virus families (Figure 7B). Sequences of *Retroviridae* and *Herpesviridae* families share the highest number of RAWs with 6 and 5 motifs, respectively. Most interestingly, the relative absent peptide ‘NGLGV’ solely appears in 156 sequences of Human coronavirus HKU1 (HCoV-HKU1) and Human coronavirus OC43 (HCoV-OC43) which both belong to the *Betacoronaviruses* genus, and tend to cause mild illness. After performing sequence alignment using Clustal Omega (58), we observed conservation of the ‘NGLGV’ motif in all 156 protein sequences of the family. More specifically, the ‘NGLGV’ pattern is localised in the S2 subunit of the coronavirus spike glycoprotein (59,60), a multifunctional protein which mediates virus invasion and fusion of the virion into host cells. We continued an *in silico* investigation on 435 full-sequenced spike glycoproteins of SARS-CoV-2 species from the current coronavirus pandemic (data retrieved on April 9, 2020 from NCBI Virus) and observed a similar (but not absent) motif ‘NGIGV’ within a similarly conserved sequence window of the same region. The replacement of human-absent ‘NGLGV’ by human-present ‘NGIGV’ suggests evolution towards host sequence mimicry and might contribute to immune evasion. Visual sequence weblogos (61) of 25-amino-acid windows around the specific RAWs (Figure 8A, 8B) demonstrate a clear amino acid consensus. Next, we extracted 71 records of full-sequenced spike glycoproteins of the same genus from various types of bat species (the latest collection date was March 2018). Although there is a varying pattern among these sequences, the most frequent motif in the examined region in bats is the ‘NGIGV’ again (Figure 8C) perhaps indicating evolutionary preadaptation. Moreover, only five sequences of spike glycoproteins from *Betacoronaviruses* in pangolins (*Manis javanica*) were available, which present a conserved ‘NGLTV’ oligopeptide within a dissimilar sequence window (Figure 8D). All protein sequences are provided in Supplementary Table 2 (Tabs 3 to 7). Although it remains unclear whether RAWs conceal underlying biological mechanisms, the fact that significant human-nullomers appear in virus sequences may provide clues to reservoir species or viral toxicity.

**Figure 8.**
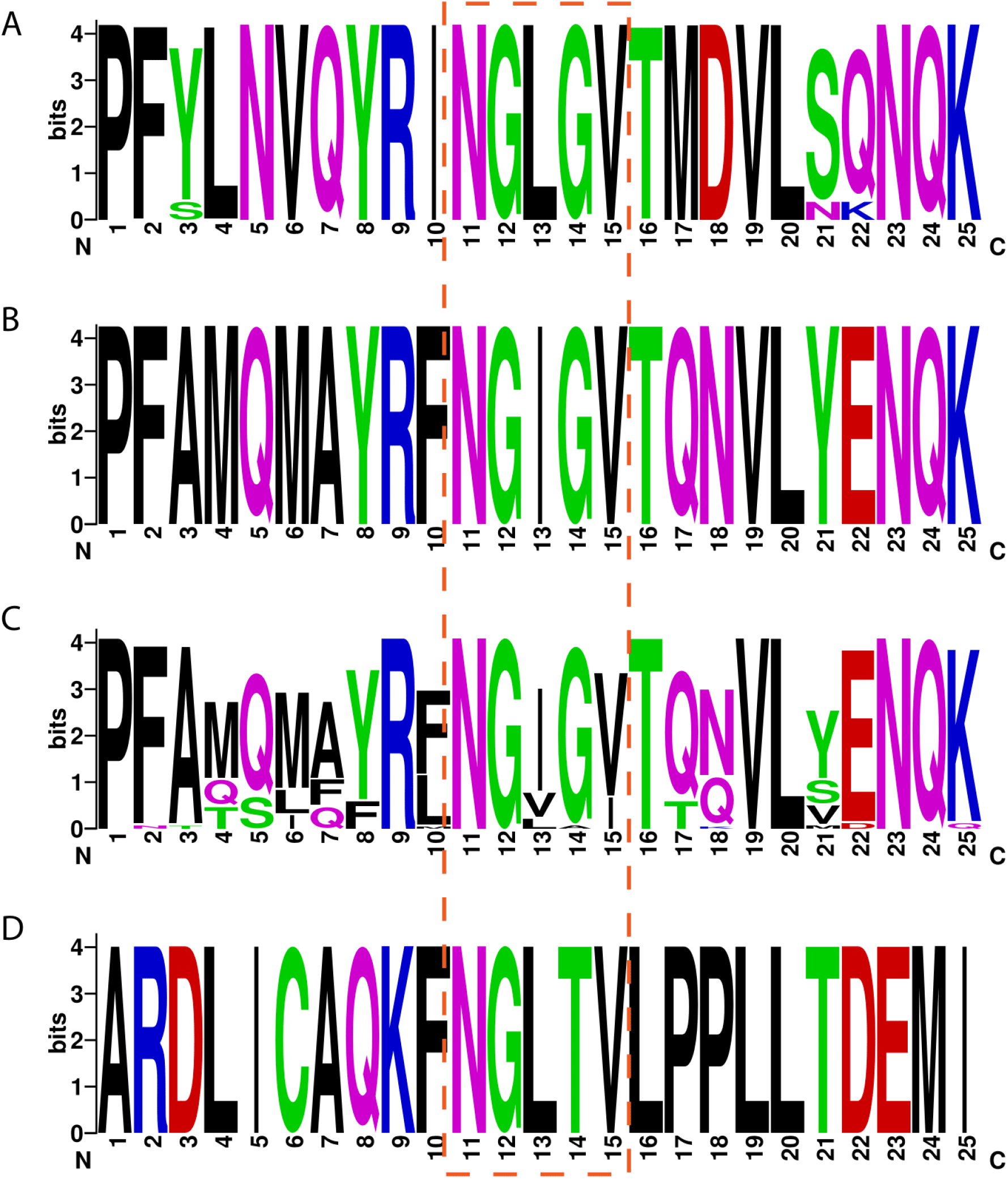
WebLogos of 25-amino-acid sequence windows from (A) 156 aligned spike glycoproteins of HCoV-HKU1 and HCoV-OC43 species in the region where the relative absent word ‘NGLGV’ occurs, (B) 435 aligned sequences from the spike glycoprotein of SARS-CoV-2 from the same protein region, 71 aligned sequence windows from spike glycoproteins of various bat species and (D) 5 aligned sequences from spike glycoproteins of Betacoronaviruses extracted from pangolins

Next, we examined whether viruses have unexpected missing motifs in their sequences. After filtering out non-complete sequences, we retrieved 147.799 individual virus genomes from NCBI Virus (data obtained in May 2020). We calculated minimal absent sequences for each virus separately and subsequently analysed them successively. More than 1.200 unique nullomers of length ranging from 4 to 10 residues were revealed significant upon Bonferroni correction at 1% cut-off. A wide species coverage is observed since the overall motifs are absent from more than 6.000 distinct species whereas, interestingly enough, the most frequent missing sequences are recognition sites of host restriction enzymes, according to the Restriction Enzyme Database (65; http://rebase.neb.com/). Restriction enzymes present in prokaryotes recognise specific sequential patterns in viruses and cleave their DNA sequences into fragments (66–67). These cutter enzymes provide a defence mechanism in host species against virus invasion and have been successfully utilized in a range of biotechnological applications and research studies (68–72). Although it is known that restriction sites confer a selective disadvantage to viruses, this has not been previously linked to nullomers. The fact that our top hits include several palindrome sequences ranging from 4 to 6 bases in length offers potential for virus nullomers to be utilised as predictors of previously unknown restriction motifs in host organisms. Furthermore, given the robustness of our evaluation method, the entire dataset can offer a valuable resource for future *in silico* research. Table 5 presents the most frequent virus nullomers which are shared between hundreds of viral species while the entire list is provided in the form of a spreadsheet (Supplementary Table 5) as well as through Nullomers Database.

**Table 5.**
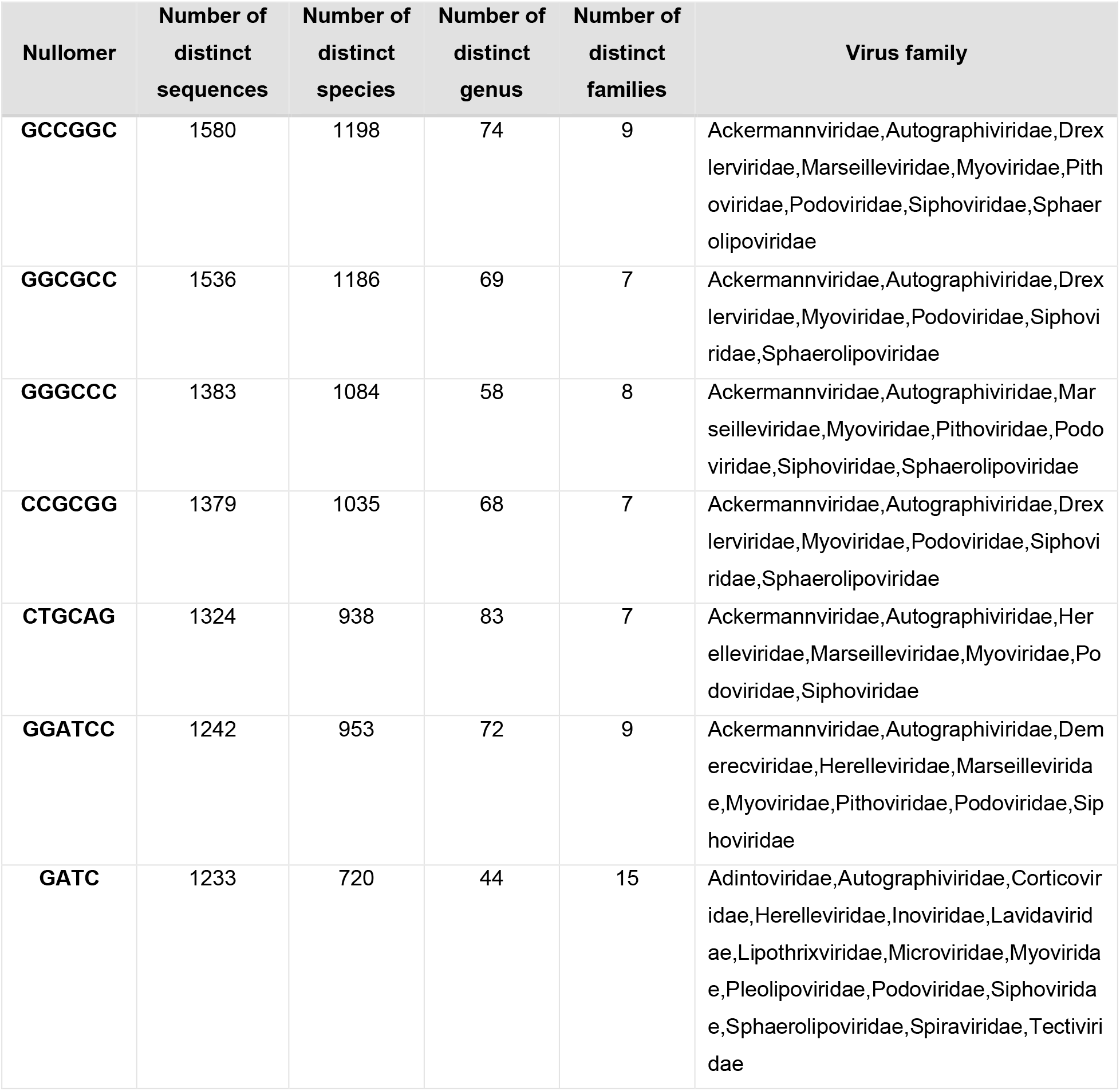
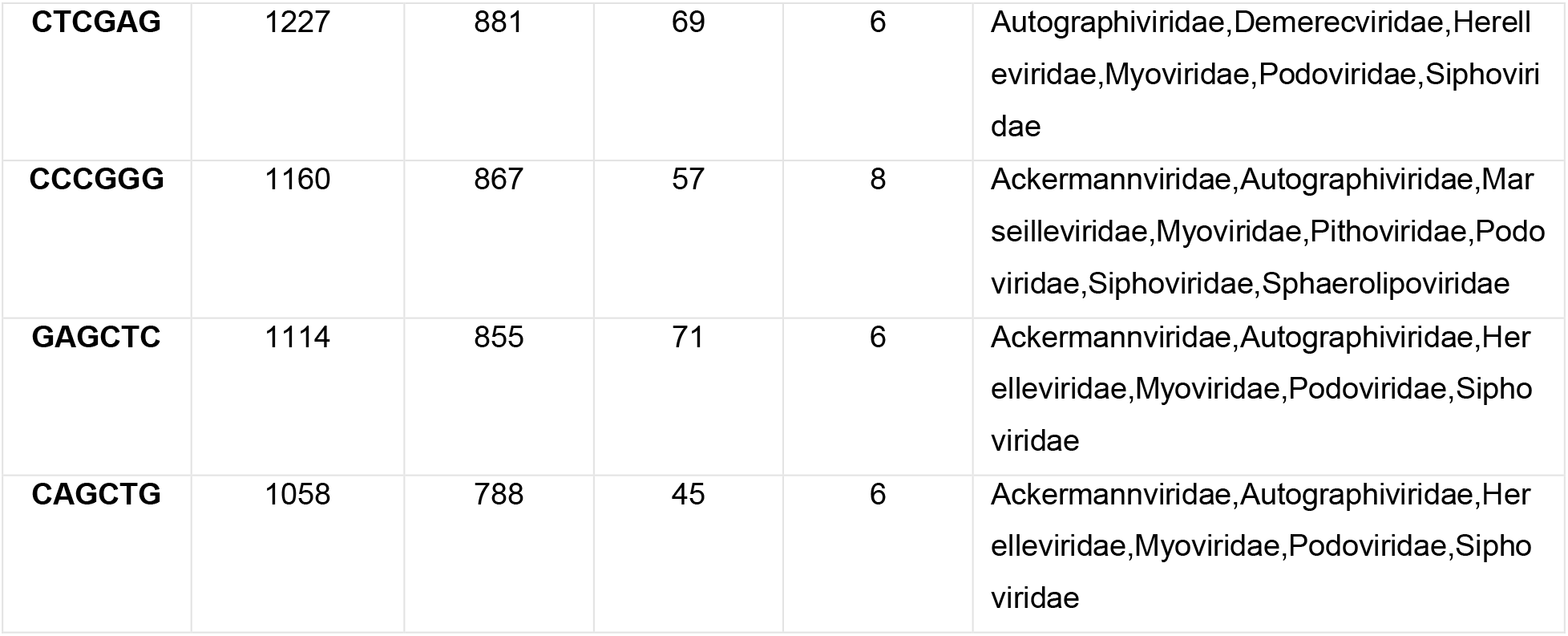
List of the most frequent (present in >1.000 distinct sequences) significant nullomers in viruses.

### Nullomers Database

To make the nullomers of the current study easily accessible and explorable, we developed an online portal available at https://www.nullomers.org. *Nullomers Database* aims to serve as a central hub of information for further investigation of minimal absent words. The provided results are significant identified genomic and peptide nullomers assessed by *Nullomers Assessor*. Main emphasis has been given to peptide nullomers and, more specifically, to regions of proteins that are prone to give rise to a significant nullomer upon a single amino acid alteration. Assuming that the deficit of the shortest absent words is governed by evolutionary factors with unfavourable consequences, then one would expect that the emergence of a nullomer upon a mutation would possibly have a negative effect. To prevent outdated information remaining in *Nullomers Database*, complex stored procedures have been developed in conjunction with an automated communication channel which retrieves information from UniProt. The gene name, protein sequence, description, sequential annotation as well as protein status (i.e. whether a protein is still active or has been deprecated and moved to UniProt Archive) are asynchronously collected from UniProt via a REST web-service. Moreover, in order to assess the significance and determine the effect of putative nullomer-making mutations, we provide functional impact annotation by Mutation Assessor (22, 23), a server which predicts functional consequences of amino acid alterations. Given the dynamic nature of the UniProt database, the information retrieval of all the above-described steps has been automated, making *Nullomers Database* a fully autonomous, scalable, and frequently updated repository. Additionally, the integration of MolArt (24), an interactive visualization plugin (Figure 9), allows for the simultaneous exploration of multiple sequential and structural features in protein nullomers. The interconnected and synchronized panels of MolArt permit users to identify co-occurrent elements in regions that are prone to engender a missing word upon a single amino acid substitution. The entire sequence annotation of the plugin is retrieved from UniProt in real-time whereas the corresponding experimental structures are dynamically fetched from Protein DataBank (or from Swiss Model Repository when a structure is not available). Therefore, the specific functionality of *Nullomers Database* ensures that the provided information can always be in line with major protein databases and automatically be enriched over time.

**Figure 9.**
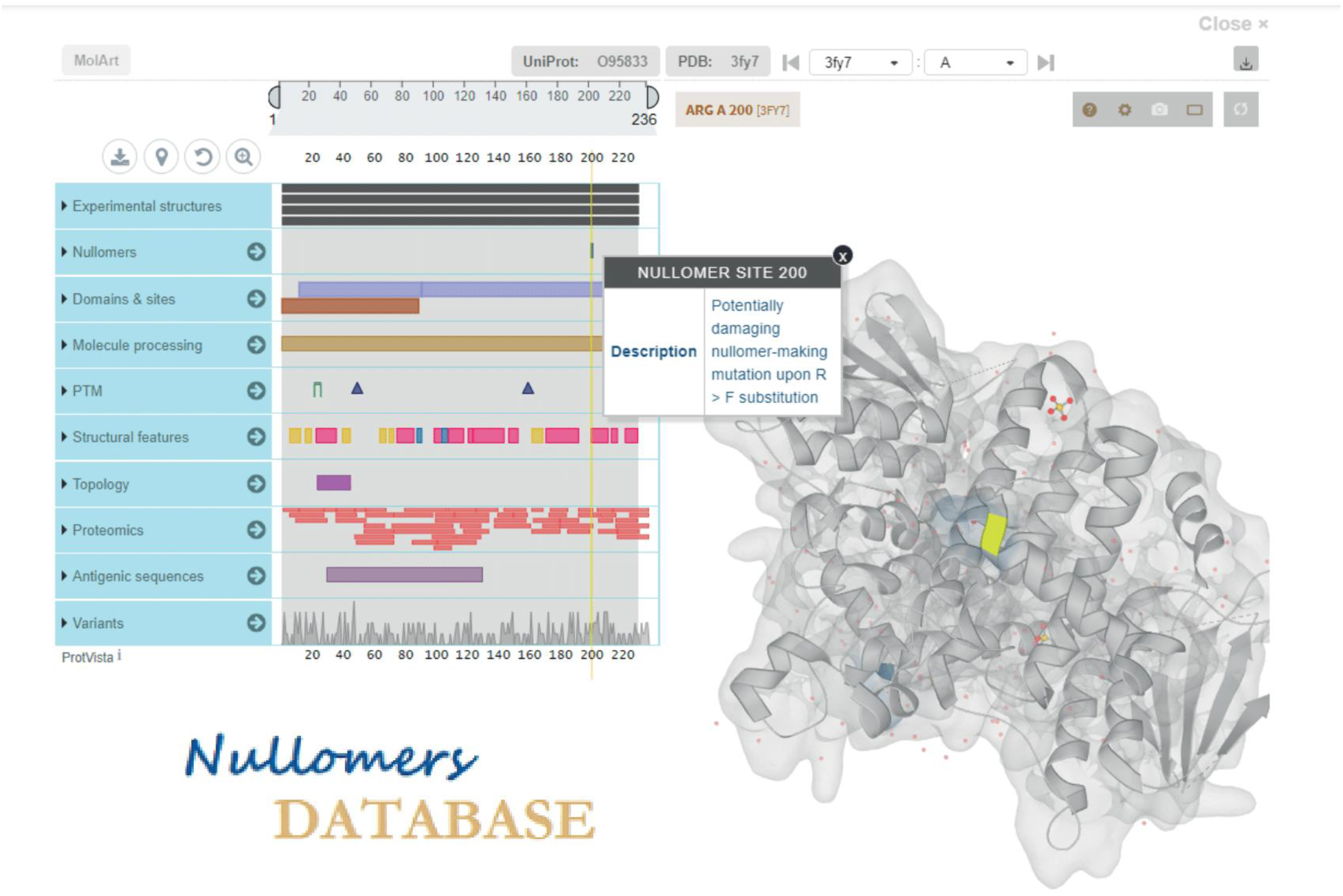
Snapshot from the graphical user interface (GUI) of Nullomers Database. Two interactive panels interconnect sequential annotation with tertiary structures offering a visual environment to explore nullomer-making mutations in proteins of interest.

## Conclusion

This paper introduces *Nullomers Assessor*, a probabilistic protocol provided as an open-source software tool, for the assessment of any set of minimal absent genomic or peptide sequences. The software offers a rigorous way of filtering missing words by Markov chains, while three statistical correction methods are available to control false positive results. We applied the script to entire genomes of hundreds of species and observed that numerous nullomers are statistically significant in multiple organisms. Moreover, we systematically examined more than 147.000 individual virus sequences and observed that the most frequent significant absent motifs are restriction recognition sites. In addition to the prevailing hypothesis that minimal absent words have gone extinct due to negative selection, we suggest that nullomers may have been replaced by more specialized sequences, which execute similar or even optimized functions.

We analysed the human and mouse proteomes and identified positions that are prone to introduce a significant missing peptide. We found that more than one fourth of human proteins can give rise to a significant nullomer upon a single amino acid substitution. We freely provide our findings in a visual, interactive, and user-friendly way via *Nullomers Database*. Taking advantage of the powerful functionalities that modern web technologies provide, we highlight protein positions which can generate a minimal absent word in their sequences.

In summary, the present study reveals significant nullomers that are unlikely to be absent by chance. Further studies should be conducted to experimentally validate and determine the actual role of nullomers as well as the extent of harmfulness behind nullomer-making mutations; hence, we anticipate that both *Nullomers Assessor* and *Nullomers Database* can be useful resources and facilitate research towards a better understanding of the still mysterious role of minimal absent words.

## Data Availability

**Project name:** Nullomers Assessor & Nullomers Database

**Project home page:** https://www.nullomers.org/

**Source code:** https://github.com/gkoulouras/nullomers-assessor

**Programming language(s):** Python, Microsoft ASP.NET, JavaScript

**Operating system(s):** Platform-independent

**Web browsers:** Google Chrome (v.76 or later), Mozilla Firefox (v.70 or later)

**License:** Apache 2.0 (http://www.apache.org/licenses/LICENSE-2.0)

## Acknowledgements

We thank and acknowledge helpful conversations with Sara Zanivan (CRUK Beatson Institute), Spyros Lytras (MRC - University of Glasgow Centre for Virus Research) and Dimitrios Vlachakis (Agricultural University of Athens). Part of this work was carried out within the financial support from the National Institute of Advanced Industrial Science and Technology.

## Author contributions’

GK and MCF contributed to the conception of the work, analysis and interpretation of the data; GK developed the scripts, designed and implemented the web database, wrote the paper and created the graphs; MCF supervised the study, wrote and proofread the final version of the manuscript. All authors read and approved the final manuscript.

## Funding

This work was supported by the Department of Information Technology and Human Factors, National Institute of Advanced Industrial Science and Technology.

## Conflict of Interest

None declared.

## Notes

### Competing Interest Statement

The authors have declared no competing interest.

## References

1. Alileche A, Hampikian G. The effect of Nullomer-derived peptides 9R, 9S1R and 124R on the NCI-60 panel and normal cell lines. BMC Cancer. 2017;17(1):533.

2. Alileche A, Goswami J, Bourland W, Davis M, Hampikian G. Nullomer derived anticancer peptides (NulloPs): differential lethal effects on normal and cancer cells in vitro. Peptides. 2012;38(2):302 – 311.

3. Goswami J, Davis MC, Andersen T, Alileche A, Hampikian G. Safeguarding forensic DNA reference samples with nullomer barcodes. J Forensic Leg Med. 2013;20(5):513 – 519.

4. Acquisti C, Poste G, Curtiss D, Kumar S. Nullomers: really a matter of natural selection?. PLoS One. 2007;2(10):e1022.

5. Barton C, Heliou A, Mouchard L, Pissis SP. Linear-time computation of minimal absent words using suffix array. BMC Bioinformatics. 2014;15(1):388.

6. Heliou A, Pissis SP, Puglisi SJ. emMAW: computing minimal absent words in external memory. Bioinformatics. 2017;33(17):2746 – 2749.

7. Pinho AJ, Ferreira PJ, Garcia SP, Rodrigues JM. On finding minimal absent words. BMC Bioinformatics. 2009;10:137.

8. Herold J, Kurtz S, Giegerich R. Efficient computation of absent words in genomic sequences. BMC Bioinformatics. 2008;9:167.

9. Silva RM, Pratas D, Castro L, Pinho AJ, Ferreira PJ. Three minimal sequences found in Ebola virus genomes and absent from human DNA. Bioinformatics. 2015;31(15):2421 – 2425.

10. Santoni D, Felici G, Vergni D. Natural vs. random protein sequences: Discovering combinatorics properties on amino acid words. J Theor Biol. 2016;391:13 – 20.

11. Chi KR. The dark side of the human genome [published correction appears in Nature. 2016 Nov 09;539(7628):318]. Nature. 2016;538(7624):275 – 277.

12. Oprea TI. Exploring the dark genome: implications for precision medicine. Mamm Genome. 2019;30(7-8):192 – 200.

13. Meyre D, Andress EJ, Sharma T, et al. Contribution of rare coding mutations in CD36 to type 2 diabetes and cardio-metabolic complications. Sci Rep. 2019;9(1):17123.

14. Chaturvedi S, Braunstein EM, Yuan X, et al. Complement activity and complement regulatory gene mutations are associated with thrombosis in APS and CAPS. Blood. 2020;135(4):239 – 251.

15. Bomba L, Walter K, Soranzo N. The impact of rare and low-frequency genetic variants in common disease. Genome Biol. 2017;18(1):77.

16. Fosgerau K, Hoffmann T. Peptide therapeutics: current status and future directions. Drug Discov Today. 2015;20(1):122 – 128.

17. Recio C, Maione F, Iqbal AJ, Mascolo N, De Feo V. The Potential Therapeutic Application of Peptides and Peptidomimetics in Cardiovascular Disease. Front Pharmacol. 2017;7:526.

18. Alexander RP, Fang G, Rozowsky J, Snyder M, Gerstein MB. Annotating non-coding regions of the genome. Nat Rev Genet. 2010;11(8):559 – 571.

19. Zhang MQ. Statistical features of human exons and their flanking regions. Hum Mol Genet. 1998;7(5):919 – 932.

20. Dai Q, Yang Y, Wang T. Markov model plus k-word distributions: a synergy that produces novel statistical measures for sequence comparison. Bioinformatics. 2008;24(20):2296 – 2302.

21. Jiang M, Anderson J, Gillespie J, Mayne M. uShuffle: a useful tool for shuffling biological sequences while preserving the k-let counts. BMC Bioinformatics. 2008;9:192.

22. Reva B, Antipin Y, Sander C. Determinants of protein function revealed by combinatorial entropy optimization. Genome Biol. 2007;8(11):R232.

23. Reva B, Antipin Y, Sander C. Predicting the functional impact of protein mutations: application to cancer genomics. Nucleic Acids Res. 2011;39(17):e118.

24. Hoksza D, Gawron P, Ostaszewski M, Schneider R. MolArt: a molecular structure annotation and visualization tool. Bioinformatics. 2018;34(23):4127 – 4128.

25. Rheinbay E, Nielsen MM, Abascal F, et al. Analyses of non-coding somatic drivers in 2,658 cancer whole genomes. Nature. 2020;578(7793):102 – 111.

26. Tamposis IA, Tsirigos KD, Theodoropoulou MC, Kontou PI, Bagos PG. Semi-supervised learning of Hidden Markov Models for biological sequence analysis. Bioinformatics. 2019;35(13):2208 – 2215.

27. Tamposis IA, Tsirigos KD, Theodoropoulou MC, et al. JUCHMME: a Java Utility for Class Hidden Markov Models and Extensions for biological sequence analysis. Bioinformatics. 2019;35(24):5309 – 5312.

28. Saw AK, Raj G, Das M, Talukdar NC, Tripathy BC, Nandi S. Alignment-free method for DNA sequence clustering using Fuzzy integral similarity. Sci Rep. 2019;9(1):3753. Published 2019 Mar 6. doi:10.1038/s41598-019-40452-6.

29. Kiesel A, Roth C, Ge W, Wess M, Meier M, Söding J. The BaMM web server for de-novo motif discovery and regulatory sequence analysis. Nucleic Acids Res. 2018;46(W1):W215 – W220.

30. UniProt Consortium. UniProt: a worldwide hub of protein knowledge. Nucleic Acids Res. 2019;47(D1):D506 – D515.

31. Di Giallonardo F, Schlub TE, Shi M, Holmes EC. Dinucleotide Composition in Animal RNA Viruses Is Shaped More by Virus Family than by Host Species. J Virol. 2017;91(8):e02381 – 16.

32. Greenbaum BD, Levine AJ, Bhanot G, Rabadan R. Patterns of evolution and host gene mimicry in influenza and other RNA viruses. PLoS Pathog. 2008;4(6):e1000079.

33. Babayan SA, Orton RJ, Streicker DG. Predicting reservoir hosts and arthropod vectors from evolutionary signatures in RNA virus genomes. Science. 2018;362(6414):577 – 580.

34. Takata MA, Gonçalves-Carneiro D, Zang TM, et al. CG dinucleotide suppression enables antiviral defence targeting non-self RNA. Nature. 2017;550(7674):124 – 127.

35. NCBI Resource Coordinators. Database resources of the National Center for Biotechnology Information. Nucleic Acids Res. 2018;46(D1):D8 – D13.

36. Hornbeck PV, Zhang B, Murray B, Kornhauser JM, Latham V, Skrzypek E. PhosphoSitePlus, 2014: mutations, PTMs and recalibrations. Nucleic Acids Res. 2015;43(Database issue):D512 – D520.

37. Creixell P, Schoof EM, Tan CS, Linding R. Mutational properties of amino acid residues: implications for evolvability of phosphorylatable residues [published correction appears in Philos Trans R Soc Lond B Biol Sci. 2012 Nov 5 ;367 (1602):3058]. Philos Trans R Soc Lond B Biol Sci. 2012;367(1602):2584 – 2593.

38. Silver DP, Livingston DM. Mechanisms of BRCA1 tumor suppression. Cancer Discov. 2012;2(8):679 – 684.

39. Foulkes WD, Shuen AY. In brief: BRCA1 and BRCA2. J Pathol. 2013;230(4):347 – 349.

40. Lazar IM, Karcini A, Ahuja S, Estrada-Palma C. Proteogenomic Analysis of Protein Sequence Alterations in Breast Cancer Cells. Sci Rep. 2019;9(1):10381.

41. Szpiech ZA, Strauli NB, White KA, et al. Prominent features of the amino acid mutation landscape in cancer. PLoS One. 2017;12(8):e0183273.

42. Al-Ssulami AM, Efficient computation of shortest absent words in complete genomes. Inf. Sci 2018; 435:59 – 68

43. Hatcher EL, Zhdanov SA, Bao Y, et al. Virus Variation Resource-improved response to emergent viral outbreaks. Nucleic Acids Res. 2017;45(D1):D482 – D490.

44. Tang D, Li B, Xu T, et al. VISDB: a manually curated database of viral integration sites in the human genome. Nucleic Acids Res. 2020;48(D1):D633 – D641.

45. Cantalupo PG, Katz JP, Pipas JM. Viral sequences in human cancer. Virology. 2018;513:208 – 216.

46. Noble WS. How does multiple testing correction work?. Nat Biotechnol. 2009;27(12):1135 – 1137.

47. Bonferroni C., Teoria Statistica Delle Classi e Calcolo Delle Probabilita, Pubblicazioni del R Istituto Superiore di Scienze Economiche e Commerciali di Firenze (Libreria Internazionale Seeber, Florence, Italy) 1936; Vol 8, pp 3–62

48. Benjamini Y, Hochberg Y. Controlling the false discovery rate: A practical and powerful approach to multiple testing. J R Stat Soc, B. 1995;57(1):289–300

49. Tarone RE. A modified Bonferroni method for discrete data. Biometrics. 1990;46(2):515 – 522.

50. Abeywickrama-Samarakoon N, Cortay JC, Sureau C, et al. Hepatitis Delta Virus histone mimicry drives the recruitment of chromatin remodelers for viral RNA replication. Nat Commun. 2020;11(1):419.

51. Venigalla SSK, Premakumar S, Janakiraman V. A possible role for autoimmunity through molecular mimicry in alphavirus mediated arthritis. Sci Rep. 2020;10(1):938.

52. Memish ZA, Perlman S, Van Kerkhove MD, Zumla A. Middle East respiratory syndrome. Lancet. 2020;395(10229):1063 – 1077.

53. Peiris JS, Yuen KY, Osterhaus AD, Stöhr K. The severe acute respiratory syndrome. N Engl J Med. 2003;349(25):2431 – 2441.

54. Ochoa D, Jarnuczak AF, Viéitez C, et al. The functional landscape of the human phosphoproteome. Nat Biotechnol. 2020;38(3):365–373.

55. Needham EJ, Parker BL, Burykin T, James DE, Humphrey SJ. Illuminating the dark phosphoproteome. Sci Signal. 2019;12(565):eaau8645.

56. Falda M, Fontana P, Barzon L, Toppo S, Lavezzo E. keeSeek: searching distant non-existing words in genomes for PCR-based applications. Bioinformatics. 2014;30(18):2662 – 2664.

57. Wu ZD, Jiang T, Su WJ. Efficient computation of shortest absent words in a genomic sequence, Inf. Process. Lett. 2010; 110:596–601

58. Madeira F, Park YM, Lee J, et al. The EMBL-EBI search and sequence analysis tools APIs in 2019. Nucleic Acids Res. 2019;47(W1):W636 – W641.

59. Li F. Structure, Function, and Evolution of Coronavirus Spike Proteins. Annu Rev Virol. 2016;3(1):237 – 261.

60. Ou X, Liu Y, Lei X, et al. Characterization of spike glycoprotein of SARS-CoV-2 on virus entry and its immune cross-reactivity with SARS-CoV. Nat Commun. 2020;11(1):1620.

61. Crooks GE, Hon G, Chandonia JM, Brenner SE. WebLogo: a sequence logo generator. Genome Res. 2004;14(6):1188 – 1190.

62. Finnegan AI, Kim S, Jin H, et al. Epigenetic engineering of yeast reveals dynamic molecular adaptation to methylation stress and genetic modulators of specific DNMT3 family members. Nucleic Acids Res. 2020;48(8):4081 – 4099.

63. Tesina P, Lessen LN, Buschauer R, et al. Molecular mechanism of translational stalling by inhibitory codon combinations and poly(A) tracts. EMBO J. 2020;39(3):e103365.

64. Zhao T, Huan Q, Sun J, et al. Impact of poly(A)-tail G-content on Arabidopsis PAB binding and their role in enhancing translational efficiency. Genome Biol. 2019;20(1):189.

65. Roberts RJ, Vincze T, Posfai J, Macelis D. REBASE--a database for DNA restriction and modification: enzymes, genes and genomes. Nucleic Acids Res. 2015;43(Database issue):D298 – D299.

66. Roberts RJ. Restriction endonucleases. CRC Crit Rev Biochem. 1976;4(2):123 – 164.

67. Sharp PM. Molecular evolution of bacteriophages: evidence of selection against the recognition sites of host restriction enzymes. Mol Biol Evol. 1986;3(1):75 – 83.

68. Arber W, Linn S. DNA modification and restriction. Annu Rev Biochem. 1969;38:467 – 500.

69. Kruger DH, Bickle TA. Bacteriophage survival: multiple mechanisms for avoiding the deoxyribonucleic acid restriction systems of their hosts. Microbiol Rev. 1983;47(3):345 – 360.

70. Ito K, Komiyama M. Site-selective scission of human genome using PNA-based artificial restriction DNA cutter. Methods Mol Biol. 2014;1050:111 – 120.

71. Lv X, Qiu K, Tu T, et al. Development of a Simple and Quick Method to Assess Base Editing in Human Cells [published online ahead of, 2020 Mar 17]. Mol Ther Nucleic Acids. 2020;20:580 – 588.

72. Schwardmann LS, Nölle V, Elleuche S. Bacterial non-specific nucleases of the phospholipase D superfamily and their biotechnological potential. Appl Microbiol Biotechnol. 2020;104(8):3293 – 3304.

73. Cheng YH, Liaw JJ, Kuo CN. REHUNT: a reliable and open source package for restriction enzyme hunting. BMC Bioinformatics. 2018;19(1):178.

74. Heberle H, Meirelles GV, da Silva FR, Telles GP, Minghim R. InteractiVenn: a web-based tool for the analysis of sets through Venn diagrams. BMC Bioinformatics. 2015;16(1):169.

75. Mun DG, Bhin J, Kim S, et al. Proteogenomic Characterization of Human Early-Onset Gastric Cancer. Cancer Cell. 2019;35(1):111 – 124.e10.

76. Lytras S, Hughes J. Synonymous Dinucleotide Usage: A Codon-Aware Metric for Quantifying Dinucleotide Representation in Viruses. Viruses. 2020;12(4):462.

